# COOBoostR: an extreme gradient boosting-based tool for robust tissue or cell-of-origin prediction of tumors

**DOI:** 10.1101/2022.10.24.513446

**Authors:** Sungmin Yang, Kyungsik Ha, Woojeung Song, Masashi Fujita, Kirsten Kübler, Paz Polak, Carolina Garcia de Alba Rivas, Patrizia Pessina, Julio Sainz de Aja, Samuel Rowbotham, Preetida Bhetariya, Antonella Dost, Aaron L. Moye, Eiso Hiyama, Hidewaki Nakagawa, Carla F. Kim, Hong-Gee Kim, Hwajin Lee

## Abstract

We here present COOBoostR (https://github.com/SWJ9385/COOBoostR), a computational method designed for the putative prediction of tissue-or cell-of-origin of various cancer types. COOBoostR leverages regional somatic mutation density information and chromatin mark features to be applied to an extreme gradient boosting-based machine-learning algorithm. COOBoostR ranks chromatin marks from various tissue and cell types which best explain the somatic mutation density landscape of any sample of interest. Through integrating either ChIP-seq based chromatin data or bulk/single cell chromatin accessibility data along with regional somatic mutation density data derived from normal cells/tissue, precancerous lesions, and cancer types, we show that COOBoostR outperforms existing random forest-based methods in prediction speed with comparable or better tissue or cell-of-origin prediction performance. In addition, our results suggest a dynamic somatic mutation accumulation at the normal tissue or cell stage which could be intertwined with the changes in open chromatin marks and enhancer sites. These results further represent chromatin marks shaping the somatic mutation landscape at the early stage of mutation accumulation, possibly even before the initiation of precancerous lesions or neoplasia.

## Introduction

Recent advances in DNA sequencing technologies have led to the development of reliable and cost-effective whole-genome sequencing methods and relevant analysis pipelines, which have been applied to numerous cancer types^1-11^, precancerous lesions^5^, normal tissues, cells and stem cells^12,13^. The somatic mutation landscape from these data revealed various genotypic alterations ranging from driver and passenger somatic mutations for cancer initiation and progression^14-17^, mutation signatures^18^, clonal and subclonal evolutions^19^, to the findings on novel structural variations including kategis^20,21^, chromothripsis^22^, and whole-genome doubling^23,24^. Although mechanisms on how these aberrations arise have not been fully defined, a number of statistical and machine-learning approaches have shown that the somatic point mutation landscapes of multiple cancer types and precancerous lesions correlated with the chromatin mark landscape^25,26^, and tissue-of-origin (TOO) or cell-of-origin (COO) predictions for different cancer types are possible by leveraging such information^27-31^.

To this day tools used for these predictions mainly utilize random forest algorithm, an ensemble learning method which uses the averaged result derived from predefined number of decision trees for the prediction. Random forest-based TOO/COO prediction algorithms, however, still face limitations such as low prediction accuracy for some cancer types, which might be due to the sparse mutation density and the requirement of a strong server-level computing power for the reasonable speed. Extreme gradient boosting (XGBoost), on the other hand, employs gradient weighing based on the prior prediction running and similarity score-based tree pruning, which in the end expected to be resistant to the unbalanced or sparse data with improved running speed. Here we describe a novel xgboost machine learning-based tool called COOBoostR, which displays a notable improvement on the prediction accuracy when there is low mutation density. It also offers the advantage of higher analysis speed in addition to the minimal requirement of computing power. To assess the validity and accuracy of the algorithm in the aspect of tissue or cell-of-origin predictions, we applied COOBoostR to the somatic point mutation landscape data from 798 tumors, 23 precancerous lesions, 35 normal stem cells, and 14 normal tissue clones/samples.

## Results

COOBoostR receives mutation and chromatin data as input after quality control (Fig. 1a) (Online Methods). We then equally divide the genomic regions into 1-megabase and calculate the values of these data in each region as has been reported previously^25^. Among the ensemble models, COOBoostR applies the XGBoost methodology to predict the TOO and/or COO. While the random forest-based TOO and COO prediction algorithms use a bagging model which performs parallel training, the boosting model of COOBoostR evaluates feature weights from sequential training. This training process improves the accuracy of the COO algorithm by strengthening models with difficult prediction. In addition, COOBoostR applies early stopping function to avoid wasting time with unnecessary repetitive training. This early stopping function is also preventing the model from over-fitting by setting the default learning rate and the number of iterations to 0.5 and 100 for sufficient training. The output of COOBoostR is a ranked list of chromatin marks, where the higher rank represents the higher similarity with the regional somatic mutation density. Eventually, the tissue or cell type corresponding to these chromatin marks is assigned as potential TOO or COO.

**Figure 1.**
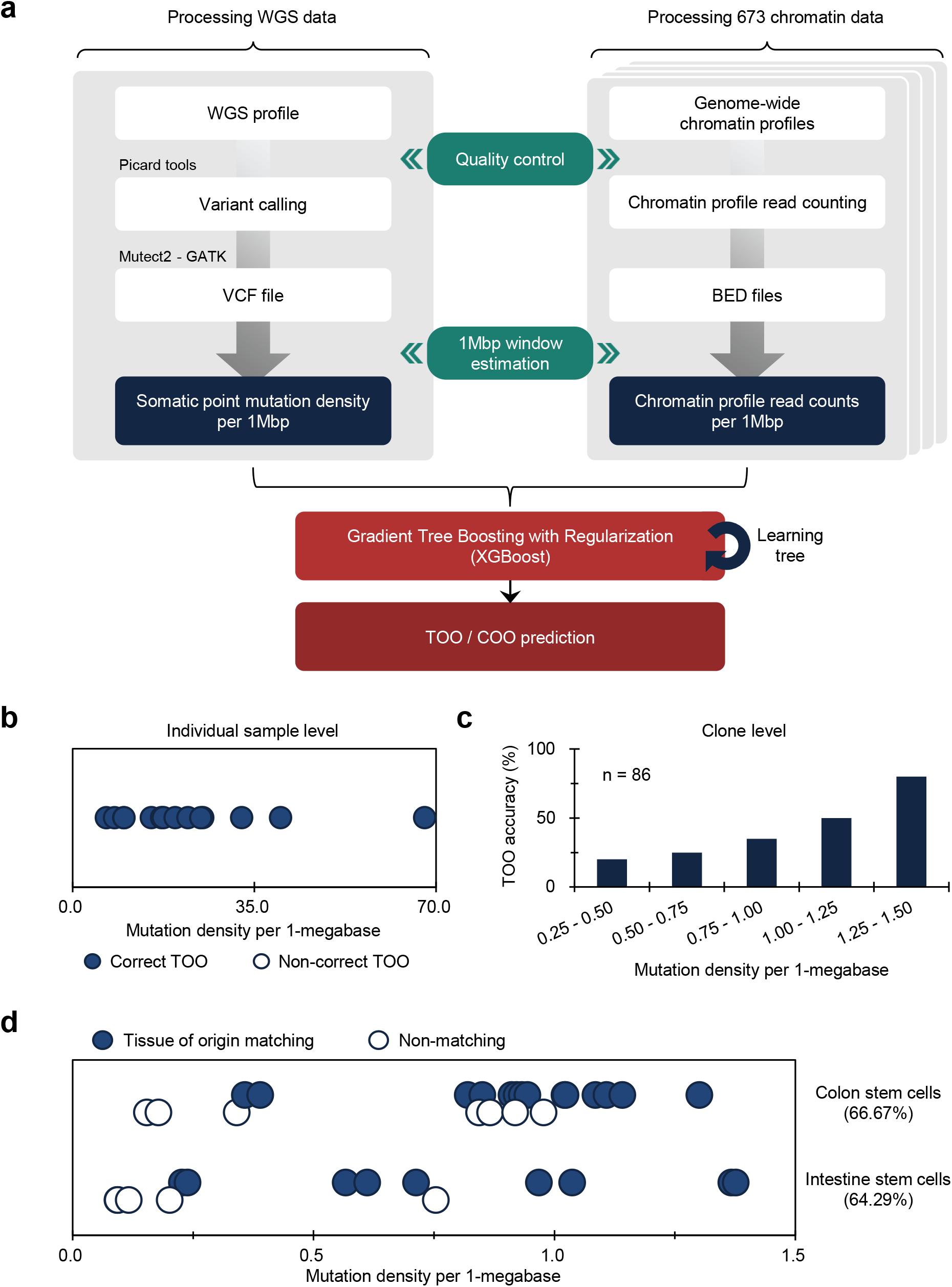
TOO/COO Prediction with COOBoostR. (**a**) COOBoostR algorithm flow diagram. A total of 870 WGS data and 673 chromatin data were subjected to the 1-megabase level preprocessing followed by XGBoost-mediated prediction stage. (**b-d**) TOO prediction accuracy for normal stem cell organoids or tissue samples harboring low mutation density (average mutation density for stem cell organoids: 0.73, liver tissue clones: 0.63). (**b**) TOO prediction accuracy for normal liver at individual sample level. Samples are aligned in order of mutation density magnitude per 1-megabase window after classifying the correctness of TOO prediction. Dots were jittered to dissect out the blue and white dots. (**c**) TOO prediction accuracy for normal liver at clone level. Histogram showing TOO prediction accuracy of liver clones with respect to the mutation density groups. (**d**) TOO prediction accuracy for colon and intestine stem cell organoids at an individual sample level. Samples matching predicted TOO are marked with solid circles, and samples that did not match are marked with empty circles. Dots were jittered to dissect out the blue and white dots. Samples are aligned in order of mutation density magnitude per 1-megabase window.

To assess speed and accuracy of COOBoostR, we compared the results from COOBoostR with a previously established random forest regression-based method^25,26^ for the prediction of tissue-of-origin for colorectal cancer, esophageal adenocarcinoma (EAC), and liver cancer. These three cancer types were previously known to display higher variance explained score compared to the other cancer types subjected to the random forest-based algorithm^25^. We were at first interested in evaluating the accuracy and reproducibility of the results after the repetitive running of COOBoostR. Our results displayed 100% reproducibility in 92.71% of tested samples (samples showing either 0% or 100% accuracy after 100 rounds of repetitions) (Supplementary Fig. 1a). We also estimated trends in TOO accuracy according to the sample mutation density per each cancer type. Among the three cancer types, 5 out of 6 colorectal cancer samples showed 100% reproducibility when the mutation density per 1-megabase of the sample was greater than 4.7, whereas the EAC and liver cancer samples exhibited 100% reproducibility of the prediction results for 100% and 95.31% of the samples regardless of the sample mutation density (Supplementary Fig. 1b). Subsequently, we conducted an accuracy comparison between the COOBoostR and the random forest-based algorithm on these samples with mutation density per 1-megabase greater than 4.7. For this, we measured the proportion of individual samples predicted as the correct tissue-of-origin with 100% reproducibility. (Supplementary Fig. 1c). In liver cancer, both algorithms performed similarly good and predicted the correct tissue-of-origin with ∼95% accuracy, whereas in colorectal and esophageal cancer, COOBoostR algorithm predicted the tissue-of-origin 1.25 and 1.05 times more samples compared to the random forest algorithm, respectively. In addition, we measured the algorithm running speed after applying COOBoostR or the random forest-based algorithm to the three cancer types. In result, COOBoostR completed the tissue-of-origin prediction process ∼389.94 times faster than the random forest-based algorithm (Supplementary Table 1). Taken together, we show that COOBoostR has reproducible results with better speed and comparable accuracy when comparing to the existing random forest-based algorithm.

Next, we tested COOBoostR to evaluate whether the regional somatic mutation data of normal tissues or cells can be best explained by the epigenome of matching normal tissues or cell types. The first dataset we used for answering such question was the whole-genome somatic mutation data from normal liver^13^. To examine the extent of the TOO matching for normal liver, COOBoostR was conducted in each individual sample from the dataset. Our results show that all 14 individual samples, with mutation density ranging from 6.6 to 67.8, were accurately matched as liver TOO (Fig. 1b). We next performed COOBoostR on individual clone-level data to see if the matching pattern could be replicated at the clonal level. Since the mutation density for the clone-level in this dataset is very low (0.80 per megabase in average per each clone), this helped us to test the performance of COOBoostR with sparse datasets, which was one of the intentions to create XGBoost-based methods^32^. For this, we matched TOO of normal liver samples at clone-level by selecting 86 clones harboring mutation density between 0.25 to 1.5 per 1-megabase out of 491 clones extracted from 14 donors. Our results show a prediction accuracy of 50% when the somatic mutation density for each clone was ∼1 per 1-megabase, and 80% if the mutation density was 1.25 or higher (Fig. 1c). These results from normal liver tissues indicate that the extent of TOO matching accuracy could depend on the mutation density of the input data.

To further test the COOBoostR-based matching prediction efficiency and to corroborate our results from the analysis using liver normal tissues and clones, we subsequently ran COOBoostR on normal adult stem cell samples which contain lower amounts of mutations (0.09∼1.38 per 1-megabase window) compared to the tumor tissues^12^. Previously it has been reported that any cancer type with low mutation density displays relatively lower TOO predictive accuracy by using the random forest-based algorithm^25,26^, we anticipated that the robustness of COOBoostR algorithm would at least partly compensate such weakness. In the case of colon adult stem cells, we divided the samples into two age-based subgroups (young (age 9 to 15) and old (age 53 to 66)) to consider the predefined age stratifications inside the cohort (Supplementary Fig. 2a). COOBoostR based TOO matching results demonstrated an accuracy of 40% for the young age subgroup samples, and 75% prediction accuracy was obtained for old age subgroup samples. In contrast, random forest-based TOO predictions resulted in 0% accuracy for the young age subgroup and 62.5% for the old age subgroup, demonstrating comparative advantage of COOBoostR over the random forest-based algorithm. This trend was consistent in the case of small intestine adult stem cells (Supplementary Fig. 2b). Again, age-based subgroups were assigned (young (age 3 to 8), mid (age 44 to 45), and old (age 70 to 87)). For the old age subgroup, both algorithms predicted correct TOO with ∼80% accuracy. However, COOBoostR-based TOO predictions for young and mid-age subgroups exhibited better performance compared to the random forest-based algorithm. While COOBoostR predicted correct TOO for 33.33% of the young age subgroup, and 100% of the mid-age subgroups, the random forest-based algorithm had an accuracy rate of 0% at the young age subgroup and 33.33% in the mid-age subgroup. Beyond an age-based subgrouping, we also checked whether there are any differences in TOO accuracy with respect to the mutation density per sample. For this, we arranged colon and small intestine adult stem cell samples according to the order of mutation density and assessed whether there was a trend. In the case of COOBoostR TOO prediction, accurate prediction was observed when the mutation density for colon adult stem cells was greater than 0.36 and the mutation density for small intestine adult stem cells was greater than 0.23 (Fig. 1d). Conversely, in the case of random forest-based TOO prediction, we observed that a relatively high mutation density is required for consistent accurate prediction (colon adult stem cell mutation density greater than 0.85, small intestine adult stem cell mutation density greater than 0.61) (Supplementary Fig. 2c). In the case of liver adult stem cells, none of the samples were predicted as the expected TOO (liver tissue), which is in line with the previous result based on the random forest-based algorithm^28^.

In addition to examining COOBoostR accuracy on the three cancer types (colorectal, EAC, and liver cancer) and normal tissue or stem cell types, we also wondered if the prediction accuracy of COOBoostR harbors comparative advantage to the random forest-based method for hepatoblastoma^10^, a pediatric neoplasm with the lowest somatic mutation density. Samples including mature (n=18) and immature (n=15) hepatoblastoma types subjected to COOBoostR had somatic mutation density of 0.133 per 1-megabase in average. When we set the liver tissue as a matching TOO for hepatoblastoma, the TOO prediction accuracy was 55.56% (5/9 samples) when the somatic mutation density was equal or higher than 0.2 per 1-megabase, whereas the accuracy was 22.22% (2/9 samples) for the same mutation density window when utilizing random forest-based algorithm (Supplementary Fig. 3). Collectively, COOBoostR displayed improved prediction accuracies for the samples harboring lower mutation density comparing to the random forest-based algorithm, albeit still showing mutation density dependent differences in accuracy performance. Also, our results are still in line with the previous finding that the TOO prediction accuracy and the mutation density are interrelated, which might be still intrinsic to the tree-based machine learning algorithm.

To further validate the accuracy of COOBoostR algorithm on the other cancer types, we conducted TOO prediction for melanoma, multiple myeloma, and glioblastoma samples (Fig. 2a)^2,6,7^. For these samples, aggregate-sample level and individual sample level predictions using random forest-based algorithm were previously reported^25^. Our analyses revealed that for melanoma samples, COOBoostR algorithm resulted in 64% accuracy for assigning correct TOO. In the case of multiple myeloma and glioblastoma samples, COOBoostR predicted correct TOOs with relatively higher accuracy (86.957% and 95.122%, respectively).

**Figure 2.**
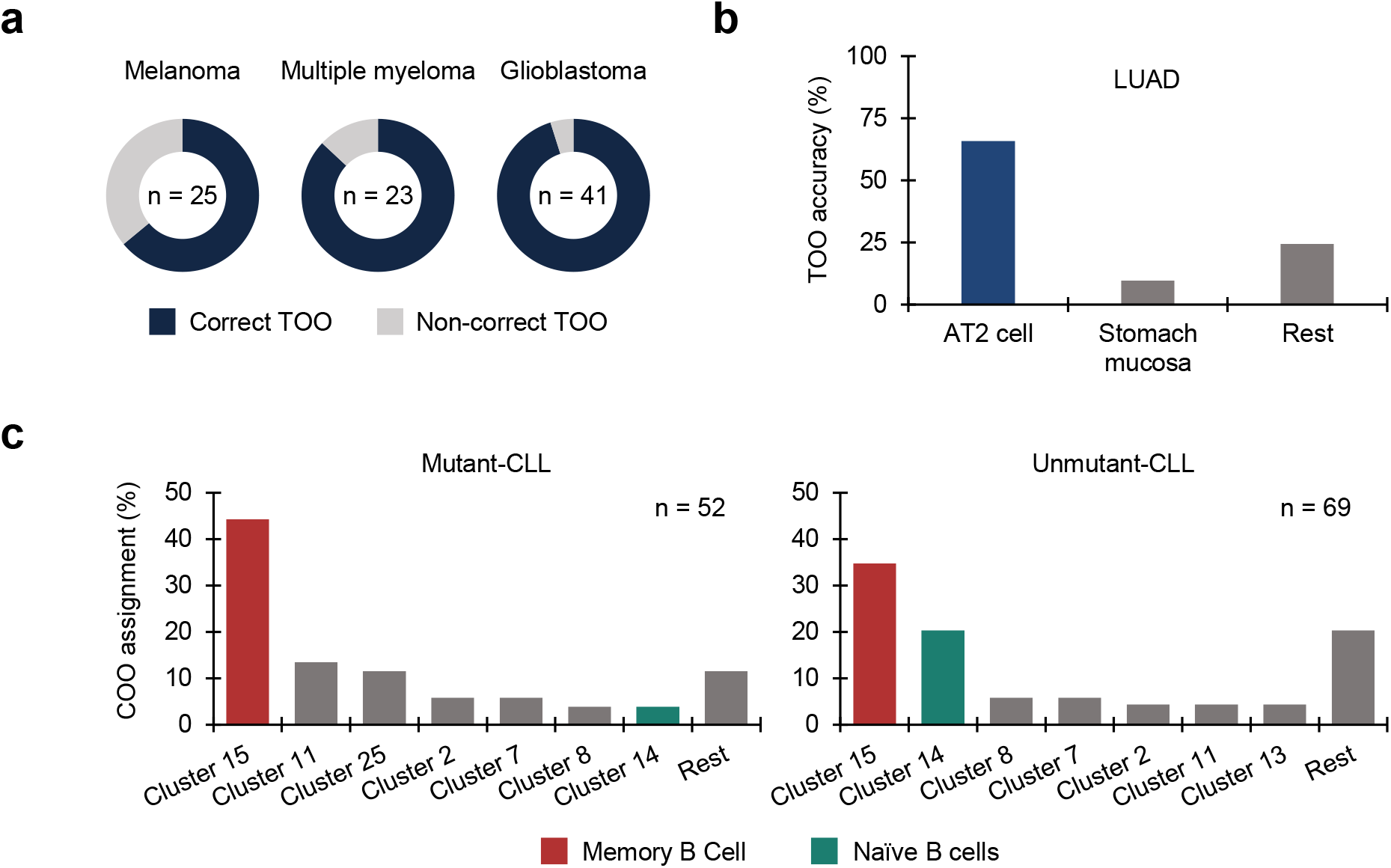
TOO/COO prediction with COOBoostR considering various input resources. (**a**) TOO prediction accuracy for Melanoma, Multiple Myeloma, and Glioblastoma. (**b)** TOO prediction accuracy for LUAD after including AT type 2 cell chromatin mark derived from bulk ATAC-seq of flow-sorted AT type 2 cells inside the chromatin data input pool. (**c**) COO prediction for IGHV-mutant and IGHV-unmutant CLL by utilizing single cell ATAC-seq data at the individual sample level. Memory B cell cluster is marked with red; naïve B cell cluster is marked with green.

In the previous multi-cancer studies, TOO/COO predictions for lung adenocarcinoma (LUAD) displayed notably lower accuracy comparing to the well-predicted cancer types like colorectal cancer and liver cancer^25^. To compensate this, we generated and applied ATAC-seq data from lung alveolar type 2 epithelial cells (AT2 cells) (Online Methods) into COOBoostR (Supplementary Table 2). We selected AT 2 cells based on the previous publications^33-35^ describing AT 2 cells as the prominent COO for LUADs in different genetically modified mouse models (GEMMs). For the ATAC-seq, we profiled a total of 4 FACS-sorted AT type 2 cell samples from 4 normal distal lung tissue donors (Online Methods). To check the data variability among individual ATAC-seq samples, we calculated the number of overlapping genes after assigning nearest genes per called peaks (Online Methods). Based on sample 4 (harboring the highest number of nearest genes assigned), 79.29% of the genes were overlapping with all the rest of the samples (Supplementary Fig. 4a). In addition, strong correlations among the individual samples were observed with respect to the regional pileup counts per 1-megabase window (Supplementary Fig. 4b). To select the best input feature format, we processed the ATAC-seq data in two ways: 1) counting the number of actual ATAC-seq peaks, or 2) calculating the number of pileup read counts per each 1-megabase window (Online Methods). As a preliminary assessment, we estimated the spearman correlation between regional somatic mutation density of LUAD^3^ at an aggregate level and the two ATAC-seq features derived from the two different data processing methods (Supplementary Fig. 4c). As a result, both the peak and the pileup-based features showed high spearman correlation values with the somatic mutation features of LUAD (spearman coefficient > 0.85) with marginal difference in rho values (difference within 0.02). Based on these pre-processing and preliminary correlation results, two ATAC-seq features of AT 2 cells were included on top of previously used chromatin mark features for running COOBoostR to obtain the TOO/COO accuracy measurement for LUAD (Fig. 2b). Our result demonstrated that the TOO prediction accuracy of LUAD increased (less than 30% to more than 60%), particularly when using the pileup-based feature is applied to COOBoostR. This results indeed underscore the need of matching TOO/COO chromatin mark feature data for better prediction accuracies and showing the utility of ATAC-seq data for the purpose of TOO/COO prediction algorithms.

So far, the chromatin feature data used in the previous TOO/COO prediction studies and the results stated in this study until this point were derived either from the whole bulk tissue, cell lines or gross level FACS-sorted cell types, which does not fully account for the detailed cell types residing in the human tissue. To overcome this, we investigated whether the single cell ATAC-seq (scATAC-seq) data could be applicable to COOBoostR. For this, we predicted the COO of chronic lymphocytic leukemia (CLL), one of the representative types of hematological malignancies with relatively high mutation load (0.86 per 1-megase per sample in our study). Since the mutation status of the immunoglobulin heavy chain gene (IgHV) is a crucial indicator not only for the different prognostic outcome in chemoimmunotherapy but also for different cell-of-origin possibilities^11,36^, we tried to perform COO prediction by classifying samples according to IgHV status (Online Methods). The analysis revealed that the cluster 15, defined as Memory B Cells^37^, was predicted to be the dominant COO in both status (Fig. 2c). However, in the case of Cluster 14 (defined as Naïve B cells), it was predicted as the second major COO only in the IgHV unmutant state. This prediction result further supports the possibility of IGHV unmutant CLL originating from the Naïve B Cells, which was proposed in previous studies^11,36^, and implicating that COOBoostR can use the scATAC-seq data as an input.

Recently, it has been suggested that the Barrett’s metaplasia (BM), a representative precancerous lesion associated with tissue metaplasia, and the EACs primarily originate from gastric cells at the level of mouse lineage tracing models^38^ and primary human samples^30,39^. To confirm whether COOBoostR prediction results align with the former outcomes, we employed Fixed-Tissue Chromatin Immunoprecipitation Sequencing (Fit-seq) data from four different tissues (BM, Squamous, Ileum, and Gastric antrum) (Online Methods), which has been previously utilized for the random forest-based TOO prediction^30^. As a first step, we tested COOBoostR algorithm on BM samples and individual matching, paired EACs (n=23). As a result, 22 out of 23 BM and EAC samples were predicted as gastric TOO, which is consistent with the gastric TOO predominancy observed by the random forest-based algorithm (Fig. 3a). Subsequently, we examined whether the prediction would change when using 1-megabase region subsets containing certain tissue-specific enhancers (Online Methods). COOBoostR predicted as gastric TOO for 12 out of 23 BM samples and 16 out of 23 EAC samples when using 1-megabase regions containing gastric tissue-specific enhancers, whereas the proportions were relatively unchanged when using 1-megabase region subsets containing squamous tissue-specific enhancers (22 out of 23 samples for BM, 23 out of 23 samples for EACs) (Fig. 3a). To examine whether these results replicate in other sample sets, we assessed the TOO prediction accuracy for the samples from two other EAC studies^4,9^. As predicted, a total of 387 out of 409 EACs^4^ and 9 out of 9 EACs^9^ were predicted as gastric TOO (Fig. 3b). In addition, the samples predicted as gastric TOO were decreased (296 out of 409 samples^4^, 4 out of 9 samples^9^) when using 1-megabse regions containing gastric tissue-specific enhancers, whereas 390 out of 409 samples^4^ and 9 out of 9 samples^9^ were predicted as gastric TOO when using 1-megabase regions containing squamous tissue-specific enhancers. These results imply that the changes in chromatin marks at or vicinity of the tissue-specific enhancer regions likely to affect the somatic mutation density profile at those regions.

**Figure 3.**
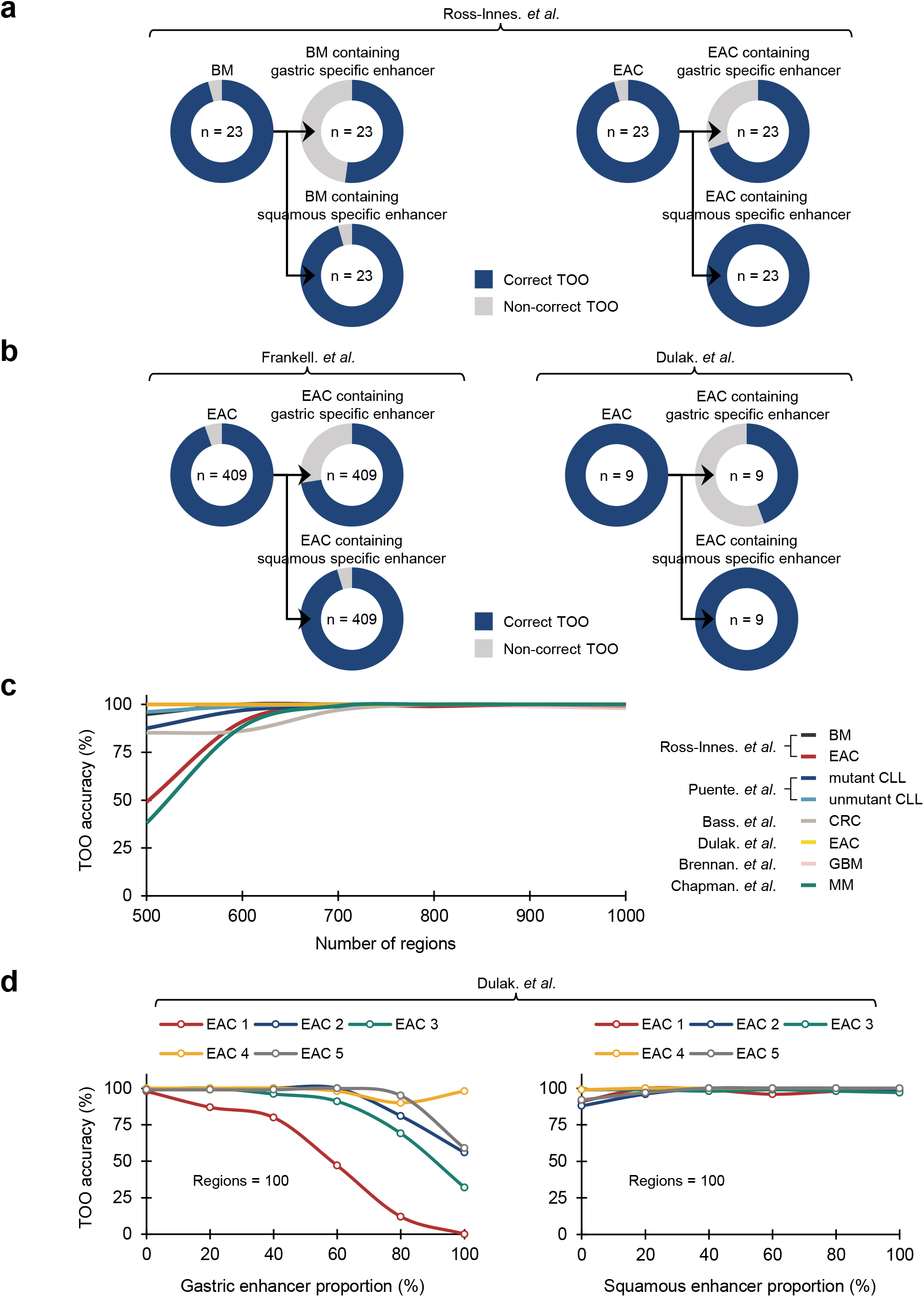
Region selection based TOO accuracy measurement for COOBoostR. (**a, b**) TOO prediction accuracy for BM and EAC derived from different study sources. In each case, TOO prediction accuracy was again measured by utilizing 1-megabase regions containing gastric specific enhancers or squamous specific enhancers. BM: Barrett’s metaplasia; EAC: Esophageal adenocarcinoma. (**c**) Region selection based COOBoostR accuracy measurement ranging from 500 to 1,000 regions for 8 sample types (7 cancer types, 1 precancerous lesion) at aggregated sample level. (**d**) Region selection analysis with respect to the portion of gastric / squamous specific enhancer containing regions for EACs at individual sample level. Enhancer inclusion ratio was varying from 0 to 100%, making up to 100 regions.

To assess any potential bias in TOO prediction results due to the differences in the number of 1-megabase regions, we conducted COOBoostR using 673 chromatin marks on 7 different cancer types and BM at aggregate sample mutation level after randomly selecting different numbers of 1-megabase region subsets (ranging from 500 to 1000, Online Methods). Cancer type dependent differences in TOO accuracies were observed only at 500 region subsets, but the prediction accuracy uniformly exceeded 97% when using 700 region subsets or more (Fig. 3c). Also, the prediction accuracies of EACs^9^ were the most consistent, reaching 100% regardless of the number of region subsets. To assess if the TOO accuracy levels are still reproduced when using different chromatin dataset, we moved onto measuring COOBoostR prediction accuracies for EACs^9^ with different region subsets utilizing the Fit-seq data. For this, we applied different region subsets according to the proportion of different enhancer containing regions (ranging from 0% to 100%, Online Methods). Results from this analysis demonstrated that the TOO prediction accuracy using 500 or 1000 region subsets was over 92% regardless of the enhancer containing region proportions and the enhancer originating tissue types (Suppmentary Fig. 5a, b). However, the gastric TOO prediction accuracies using 100 region subsets mostly decreased as the proportion of the gastric enhancer containing regions become higher, whereas the prediction accuracy using 100 region subsets with different squamous enhancer containing region proportions was consistently high (over 88%) (Fig 3d). These results provide evidence of minimal bias towards the number of 1-megabase regions on COOBoostR prediction and reinforce the role of tissue specific chromatin mark profiles on the somatic mutation accumulation during the course of precancerous lesions and cancer development.

## Discussion

COOBoostR is based on the extreme gradient boosting method (XGBoost)^32^, which was originally designed to cope with the issues raised by tree-based learning approaches (relatively low throughput with high consumption of computing power, poor prediction accuracies when using sparse data matrix, etc.). Based on the gradient weighing and tree pruning process embedded in the algorithm, we utilized the XGBoost methodology to select and rank tissue or cell-level chromatin mark features with respect to the relationship with somatic point mutation density of normal tissue / pre-cancerous lesions / tumor samples with improved speed and computing power comparing to the existing random-forest based algorithm. Leveraging this machine learning based feature ranking, TOO/COO predictions on samples harboring diverse mutation densities were performed by incorporating different chromatin mark input datasets.

Although there are several publications which described strong correlative measurements between TOO/COO chromatin marks and somatic mutation landscape of tumors or precancerous lesions^25,26,28,31,40^, no pre-existing reports tackled into the question on whether the regional somatic mutation density profiles of normal tissues or cells indeed best explained by the matching TOO or COO mark profiles. We postulated that the accurate matching of TOO/COO for normal tissues or cell types through utilizing our algorithm with somatic mutation inputs would not only confirm that COOBoostR is working as designed but also support the argument on the role of chromatin marks shaping somatic mutation landscape and utilizing such concept for predicting TOO/COO for various cancer types. Albeit revealing limitations possibly due to the very low mutation density for some of the samples, COOBoostR showed that the regional somatic mutation data of normal tissues or cells can be best explained by the epigenome of matching normal tissues or cell types, which is more apparent comparing to the random forest-based algorithm for the samples harboring lower mutation density. In line with this, similar phenomenon was observed for the hepatoblastoma samples, which showed the lowest mutational density among human cancers^10^. It would be worth investigating to see if this pattern applies to other normal tissues, cell types and pediatric tumors.

COOBoostR was able to employ diverse types of chromatin data inputs, ranging from ChIP-seq, Repli-Seq, DNase-seq, Fit-seq (which have also been utilized in the previous studies based on random forest-based algorithms) to bulk ATAC-seq and scATAC-seq, which has not been used in any of the previous studies related to the TOO/COO predictions. For the bulk ATAC-seq, we generated the data from FACS-sorted AT 2 cells and postulated that the most crucial reason for the low prediction accuracy displayed in previous studies^25^ is due the absence of relevant TOO/COO markers related to the lung tissue or cell types as the input data. Also, we utilized the scATAC-seq data encompassing hematopoietic cell types to cover major limitations on the cell type coverage for hematopoietic cancer types. When using these two datasets, COOBoostR results do show consistency with the COO of LUAD and CLL which has been proposed in several previous publications^11,33,36^. Utilizing scATAC-seq data from other normal tissues to directly predict putative COO for various cancer types with unknown or controversial COO would be an important next step for future research.

In this study, we examined whether COOBoostR prediction results per individual sample would change when utilizing 1-megabase region subsets containing tissue-specific enhancers. This analysis was based on the hypothesis that the chromatin marks at or proximal to the TOO-specific enhancer regions would change during the progression to BM mainly by the loss of such enhancers, and somatic mutation accumulation patterns inside those regions would change dynamically during the progression to BM, thus less correlating with the original TOO chromatin marks and ultimately affect COOBoostR accuracy. In line with our hypothesis, marked decrease of gastric TOO-predicted samples for BM and EAC were observed when using 1-megabase regions containing gastric tissue-specific enhancers, whereas the proportions of gastric TOO-predicted samples were conserved when using 1-megabase region subsets containing squamous tissue-specific enhancers. This phenomenon was recapitulated for the samples from two other studies by demonstrating reduced TOO prediction accuracy when using 1-megabase regions containing gastric tissue-specific enhancers. This phenomenon was more evident when the number of region subsets became lower. Although the window size of the enhancer regions is far less than 1-megabase, our results do emphasize the changes in tissue specific chromatin marks during precancerous lesions/malignant tumor progression, at least in the case of BM and EAC, is sufficient to be captured as COOBoostR prediction accuracy changes.

In conclusion, COOBoostR showed at least comparable or better TOO/COO prediction speed and accuracy with wider options of chromatin data inputs comparing to the random forest-based algorithm. Also, our results provide several lines of evidence for dynamic somatic mutation accumulation at the normal tissue or cell stage which could be intertwined with the changes in open chromatin marks and enhancer sites. COOBoostR would facilitate cancer COO investigations and predictions in human, which would be critical to early cancer diagnosis and selecting treatment options.

## Online Methods

### Somatic mutation data derived from whole-genome sequencing

We calculated regional somatic mutation density for 870 individual cancer genomes derived from several cancer types, precancerous lesions, normal stem cells, and normal tissues and cells. The International Cancer Genome Consortium (ICGC), which include Pan-Cancer Analysis of Whole genomes (PCAWG), Accelerating Research in Genomic Oncology (ARGO), and European Genome-Phenome Archive (EGA), has granted permission to use 64 liver cancer genomes (LIV)^1^, 41 glioblastoma genomes (GBM)^2^, 41 lung adenocarcinoma genomes (LUAD)^3^, 413 esophageal adenocarcinoma genomes (EAC, Frankell. *et al*)^4^, and 23 pairs of Barrett’s metaplasia (BM) matching with esophageal adenocarcinoma genomes (EAC, Ross-Innes. *et al*.)^5^. In our study, Barrett’s metaplasia genomes were employed as a representative case of precancerous lesions. In the case of 413 EACs deposited in ICGC ARGO, four samples (DO234285, DO234363, DO234413, DO234462) with hypermutations were excluded from TOO/COO predictions and the genomic coordinates of the variants for each sample were converted from Genome Reference Consortium Human Build 38 (GRCh38) to Genome Reference Consortium Human Build 37 (GRCh37) using CrossMap^41^. From The database of Genotypes and Phenotypes (dbGaP), we have been granted authority for data use of 25 melanoma genomes (MEL)^6^, 23 multiple myeloma genomes (MM)^7^, 9 colorectal cancer genomes (CRC)^8^, and 9 esophageal adenocarcinoma genomes (EAC, Dulak. *et al*.)^9^. We have been also granted access to published 33 hepatoblastoma genomes from Hiroshima University and RIKEN^10^. In the case of chronic lymphocytic leukemia (CLL), which consists of 52 IGHV-mutant genomes and 69 IGHV-unmutant genomes, somatic mutation data were extracted from Supplementary Table 2 of the publication^11^. For normal stem cells, we gathered tissue-specific somatic mutation accumulation in adult stem cells from publicly available datasets^12^, which include 21 colon stem cell genomes and 14 intestine stem cell genomes. As a representative case of normal tissue, we not only extracted 14 normal liver genomes, but also 491 normal liver clone genomes within samples from published data source^13^. Since the mutation rate of normal liver clone genomes per 1-megabase window were distributed from 0.25 to 1.5, we divided each interval into 0.25 mutation rate units and randomly selected a total of 86 clonal samples from each interval. In order to measure the regional mutation density for each sample, autosomes were split into 1-megabase regions excluding areas related to centromeres, telomeres, and low quality unique mappable base pairs. After that, we aggregated the frequency of variations in each 1-megabase region and established somatic mutation profile for individual samples. This mutation counting process was carried out using BEDOPS^42^, and based on the Genome Reference Consortium Human Build 37 (GRCh37).

### Chromatin data derived from Chromatin Immunoprecipitation sequencing (ChIP-seq)

A total of 673 ChIP-seq data, which include human primary tissues and cell lines, were extracted from ENCODE^43^, IHEC^44^, and the NIH Roadmap Epigenomics Consortium (release 9)^45^. In our study, these ChIP-seq features were classified as a total of 132 tissue or cell types according to the original source^31^. Moreover, the ChIP-seq data utilized in this study contain two kinds of repressive histone modifications (H3K27me3 and H3K9me3) and five kinds of active histone modifications (H3K27ac, H3K36me3, H3K4me1, H3K4me3, and H3K9ac). Consistent with the mutation profile counting, we calculated ChIP-seq reads count in each 1-megabase region based on the human genome version GRCh37 (hg19).

### Bulk and single-cell chromatin accessibility data derived from Assay for Transposase-Accessible Chromatin sequencing

On top of the epigenetic mark feature datasets described above, ATAC-seq data from lung AT type 2 cells was newly generated in our study specifically for the purpose of our COO prediction study. As a first step, distal lung coming from 4 different donors (Lu8, Lu13, Lu14 and Lu20) were dissociated with liberase and sorted for EPCAM+ (biolegend PE/Cy7 anti-human CD326 (Ep-CAM) [9C4] 100 tests cat. 324222), CD31-(biolegend PE anti-human CD31 [WM59] 100 tests cat 303106), CD45-(biolegend PE anti-human CD45 [2D1] 100 tests cat 368510), NGFR-(biolegend APC anti-human CD271 (NGFR) [ME20.4] 100 tests cat. 344108) and HTII-280+ (Terrace biotech HTII-280 mouse IgM cat. TB-27AHT2-280). 50 × 10^^3^ EPCAM+CD45-CD31-NGFR-HTII-280+ cells were sorted per donor. DNA extraction, transposition and amplification following previous methods^46^ from the Howard Chang lab. 80nM final concentration of transposase was used. Libraries of donors were pooled together on a total concentration of 17.7 ng/ul, validated the quality through Tapestation (High sensitivity D1000 Screen Tape) and used Nextseq (mid output).

ATAC-seq analysis was performed using the bcbio-nextgen toolkit (via the ChIP/ATAC-seq pipeline) (1). The guidelines for ATAC-seq analysis provided by ENCODE was followed as best practice (2). Quality control (QC) metrics were based upon recommendations from Yiwei Niu (3). Reads were filtered and trimmed with Atropos (4)^47^. High quality reads were mapped to the human genome (build GRcH37/hg19) using Bowtie2 (5)^48^. After filtering reads from mitochondrial DNA, properly paired reads with high mapping quality (MAPQ score >10, non-duplicates, uniquely mapped) were retained for further analysis. ATAC-seq specific QC reports were generated using ataqv which produces a web-based visualization and analysis report containing basic QC metrics (6). The functions ‘alignmentSieve’ and bamPEFragmentSizefunction’ from Deeptools (7)^49^ and Samtools (8)^50^ were used to isolate fragments in nucleosome free regions (NFRs), mono-nucleosome and di-nucleosome regions from aligned BAM. Accessible regions (ATAC-seq peaks) were called from BAM files using the MACS2 algorithm (9)^51^ followed by a quality check with the ChIPQC Bioconductor package (10)^52^. The bigwig files (bin size=20) were visualized using IGV (11)^53^. After peak calling, post-processing of the data was conducted by aggregating pileup counts with respect to each 1-megabase window region per sample, then calculate the average value of the 1-megabase region pileup counts. The pileup sequences at the boundary of 1-megabase regions were assigned to the region which harbors more part of the actual sequence. In order to estimate the extent of peak overlap among the ATAC-seq data from each individual sample, we first performed nearest gene assignment for each annotated peak using HOMER’s annotatePeaks.pl software^54^. Subsequently, we confirmed the number of matching genes among the ATAC-seq samples as a representative of how much the peaks were overlapping in the form of a Venn diagram. (Supplementary Fig. 4a)

To predict COO for hematopoietic cancers, single-cell ATAC-seq (scATAC-seq) data representing diverse hematopoietic cells^37^ were subjected to the analysis. Similar to the bulk-ATAC-seq data, post-processing of the data was conducted by adding the pileup counts with respect to each 1-megabase window region per cell, then calculate the multiple-cell aggregate value of the 1-megabase region pileup counts with respect to each cell type. The pileup sequences at the boundary of 1-megabase regions were assigned to the region which harbors more part of the actual sequence. For the count data from LSCs and Blasts, we further averaged the count data for two individuals.

### COOBoostR: XGBoost-based feature selection algorithm for TOO/COO prediction

Identification of putative TOO/COO for cancer types can be crucial for decoding the responsible tissue or cell types subjected for monitoring-based cancer prevention and understanding the somatic mutation accumulation mechanisms intertwined with the cancer development. Pre-existing methods for TOO/COO identifications include lineage-tracing mouse models and organoids, which are either heavily depending on a particular genetically engineered mouse models or time-consuming with extremely low throughput. Bioinformatics and machine learning based approach to predict TOO/COO for human tumor samples is unique in a sense that this is the sole practical method for estimating the TOO/COO directly for human samples with a reasonable throughput.

XGBoost is an extendible and state-of-the-art algorithm of gradient boosting machines which has proven to push the limits of computing power for boosted trees algorithms^32^. It was developed for the sole purpose of model performance and computational speed. Boosting is an ensemble technique in which new models are added to adjust for existing model errors. Model is recursively added until the error is no longer small and is adjusted according to the hyperparameter. Furthermore, gradient booting is an algorithm that combines the previous model with the new model that predicts the residuals of the previous model to make the final prediction, which then updates the weights of the model. To minimize the loss when adding new models, gradient descent algorithm is utilized. Also, the performance was significantly improved by using multiple cores of a CPU and reducing the lookup times of individual trees created in the XGBoost.

COOBoostR was created to achieve higher accuracy and speed comparing to the current state-of-art random forest-based algorithm by taking advantage of the XGBoost algorithm. The input of COOBoostR is a matrix containing regional somatic mutation density for each 1-megabase window (2,128 rows) of the human genome, whereas the output provides top20-ranked chromatin marks derived from responsible tissue or cell types which show high correlations to the regional somatic mutation densities. For this algorithm to work accordingly, we applied backward elimination manners to seek a minimal set of predictors for each genome. Also, we trained COOBoostR model with 10-fold cross-validation on the complete set of variables and determined the importance of all the variables in the model. We then ranked the predictors according to their importance and determined the top 20 variables. The most important variable (top 1) is obtained through the training 20 models and the backward elimination method. The last predictor variable represents the most similar landscape regarding the 1-megabase level regional mutation density.

Tuning COOBoostR can involve many hyperparameters including “n estimators”, which determines the epoch of the model, “early stoppong rounds” functions to check for overfitting, “learning rate”, gamma”, “max depth”, etc. To customize the model, hyperparameter tuning is possible. For example, several learning rate variable options (0.05, 0.1, 0.3, 0.4, 0.5, 0.6, 0.7, 0.8, 1.0) are available, which 0.5 was selected as a default value. In addition, default value of 20 was set for n estimators variable, and gamma of 0 (0, 0.5, 1, and 2 can be options) was set as a default. Finally, max depth of 6 was used as the default value among 1, 3, 6, and 8.

### Random Forest-based TOO/COO prediction

Our TOO/COO predictive analysis based on random forest regression was performed by reflecting and modifying previous research^25,26,28^. Once the training sets of each tree were constructed, the mean squared errors were then measured from out-of-bag data to determine the importance of each variable. In each tree, the values of these variable were randomly permutated and estimated. The raw importance value of variable m was calculated by subtracting the mean squared error between the untouched out-of-bag data and the variable-m-permuted data. Consequently, the importance ranking of each variable was estimated from the average score of the variable m in the entire tree. To predict regional mutation density for each sample, we generated 1000 random forest trees based on 673 epigenetic features. In addition, the TOO/COO for each sample was predicted from the tissue/cell type of top 1 epigenetics marker identified by employing the greedy backward elimination method. The random forest models at each stage were repeatedly tested 1000 times. We employed these random forest based TOO/COO prediction algorithms to compare the speed and accuracy with COOBoostR algorithm. For the speed comparison between the two algorithms, the genomes of liver cancer, esophageal adenocarcinoma (Ross-Innes. *et al*.), and colorectal cancer was employed. In comparing the prediction accuracy of the two algorithms, not only these cancer type samples, but also normal adult stem cell genomes were employed.

### COOBoostR and region subset-based analysis

Aggregated regional somatic mutation density for 7 cancer types (chronic lymphocytic leukemia – IGHV mutant type (mutant CLL), chronic lymphocytic leukemia – IGHV unmutant type (unmutant CLL), esophageal adenocarcinoma from Ross-Innes. et. al (EAC), esophageal adenocarcinoma from Dulak. et. al (EAC), colorectal cancer (CRC), multiple myeloma (MM), and Glioblastoma (GBM)) and 1 pre-cancerous lesions (Barrett’s Esophagus (BE)) were subjected to the analyses. For generation of aggregated regional somatic mutation density matrix, the number of somatic mutations per 1-megabase region was added up for all of the samples corresponding to each cancer type. Also, individual sample-level regional somatic mutation density for EAC (5 samples) cancer type was utilized for the analyses. These individual samples were selected based on their COOBoostR prediction results using entire regions matching to the TOO/COO prediction performed by using aggregated regional somatic mutation density of the corresponding cancer type.

For aggregated regional somatic mutation density data, COOBoostR was performed on 6 different region subset cases (500, 600, 700, 800, 900, and 1000 regions), and the accuracy (defined by predicted TOO/COO when using all of the 2,128 regions) calculated by 100 random sampling iterations was derived for each case. For individual sample-level regional somatic mutation density data, subset cases consisted of a total of 21 different number of regions (50, 100, 150, 200, 250, 300, 350, 400, 450, 500, 550, 600, 700, 800, 900, 1000, 1200, 1400, 1600, 1800, and 2000 regions), and the accuracy measurements were derived by 20 times of random region subsampling.

### Tissue specific enhancer containing region subset based TOO predictions

For the region selecting-based TOO prediction of BM and EACs containing tissue specific enhancers, Fixed-Tissue Chromatin Immunoprecipitation Sequencing (Fit-seq) data derived from 4 different tissues (Gastric, Barrett’s, Squamous, Ileum) were employed^30^. We first selected the regions containing tissue specific enhancer regions (gastric or squamous) from a total of 2128 regions. Then, among the four types of Fit-seq markers, samples predicted as a gastric tissue mark were considered as the accurate TOO predicted samples for both BM and EACs. In Figure 3a and b, we measured the accuracy of TOO prediction for BM and EACs at the individual sample level. In Figure 3d and Supplementary Fig. 5, TOO accuracy was measured after subsetting regions based on the proportion of tissue specific enhancer containing regions (0, 25, 50, 75 to 100% for 5 EACs (covering the lowest to the highest somatic mutation density within the sample set). Specifically, random selection of the enhancer containing regions is first performed with respect to the predefined proportion of the enhancer containing regions, then the random selection of the other regions is conducted to fill the total number of the intended subset regions if there is any room left. A total of 100 rounds were executed per each sample and the fixed region number, then the TOO prediction accuracy was calculated.

## Code availability

Source code for COOBoostR is available at https://github.com/SWJ9385/COOBoostR

## Acknowledgement

We thank Dr. Ramesh A Shivdasani and Harshabad Singh for providing Fixed-Tissue Chromatin Immunoprecipitation Sequencing data with helpful comments.

K.K. is supported by the Private Excellence Initiative Johanna Quandt of the Stiftung Charité at BIH.

A.L.M. is supported by a Damon Runyon Cancer Research Foundation postdoctoral fellowship (no. DRG:2368-19) and a Postdoctoral Enrichment Program Award from the Burroughs Wellcome Fund (no. 1019903). This work was supported by R35HL150876, the Thoracic Foundation (C.F.K.).

## Contributions

H.L., K.H., H.K. and C.F.K. contributed for conceiving and designing the analysis and the experiments. M.F., P.P., J.A.S., C.G.A.R., P.P., S.R., P.B., D.A., A.L.M., E.H. and H.N. contributed reagents, materials, analysis tools or the data. S.Y., K.H., H.L., W.S. and B.P. analyzed and interpreted the data. H.L., K.H., S.Y., W.S., J.A.S., C.G.A.R. and P.B. wrote the paper. K.K. provided scientific insight, contributed to the interpretation of the data and edited the manuscript.

