## Supplementary Figures and Tables for "COOBoostR: an extreme gradient boosting-based tool for robust tissue or cell-of-origin prediction of tumors"

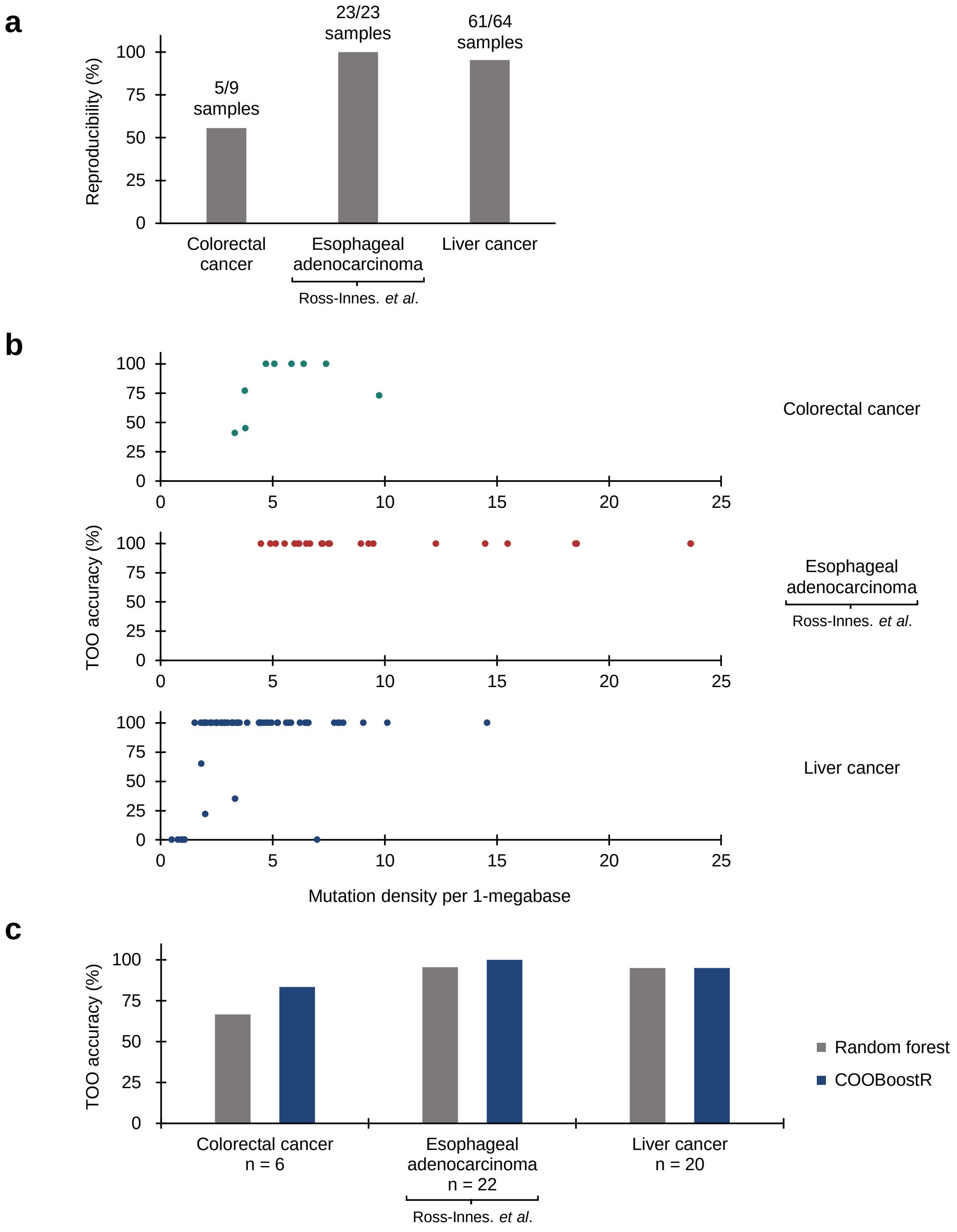


**Supplemental figure 1.** TOO accuracy comparisons between COOBoostR and Random forest-based algorithm. (**a**) Reproducibility of COOBoostR. The number of samples showing either 0% or 100% accuracy is represented above the bar graph. (**b**) TOO prediction accuracy for colorectal cancer, esophageal adenocarcinoma, and liver cancer at an individual sample level using 100 repeated COOBoostR algorithm. Samples are aligned in order of mutation density magnitude per 1-megabase window. (**c**) TOO accuracy comparison between COOBoostR and random forest-based algorithm.

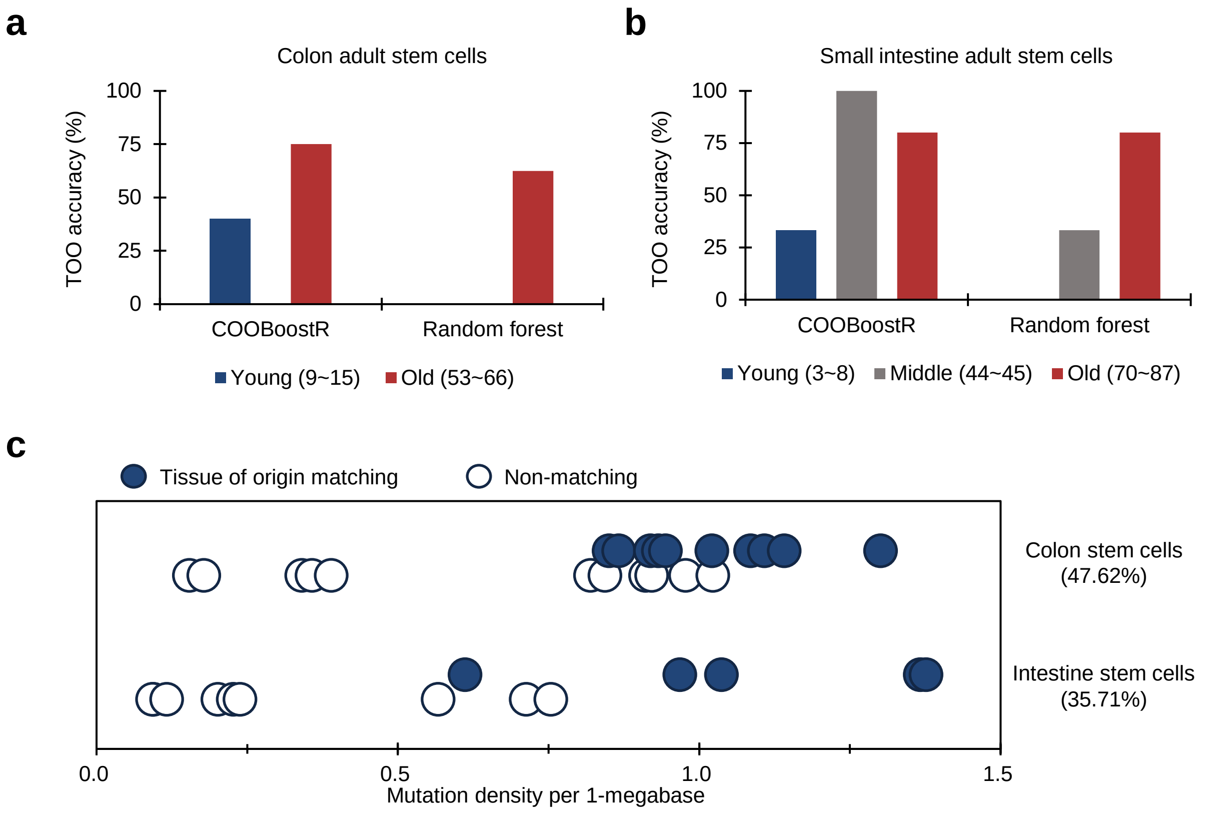


**Supplemental figure 2.** TOO accuracy comparison for normal adult stem cells between COOBoostR and Random forest-based algorithm. (**a**) TOO accuracy comparison for colon adult stem cells according to age-based subgrouping. (**b**) TOO accuracy comparison between for small intestine adult stem cells according to age-based subgrouping. (**c**) TOO prediction accuracy for colon and intestine stem cell organoids at an individual sample level using random-forest algorithm. Samples matching predicted TOO are marked with solid circles, and samples that did not match are marked with empty circles. Dots were jittered to dissect out the blue and white dots. Samples are aligned in order of mutation density magnitude per 1-megabase window.

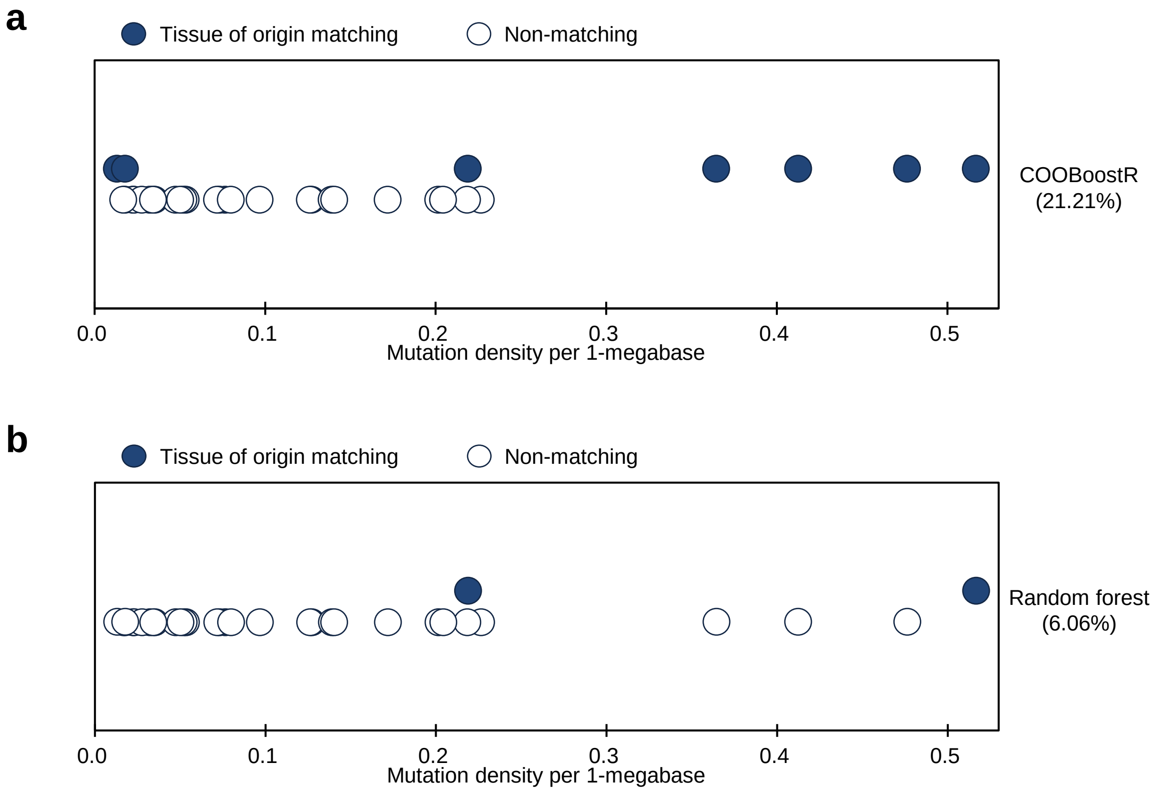


**Supplemental figure 3**. TOO prediction accuracy for hepatoblastoma at an individual sample level using (**a**) COOBoostR and (**b**) Random forest-based algorithm. Samples matching predicted TOO are marked with solid circles, and samples that did not match are marked with empty circles. Dots were jittered to dissect out the blue and white dots. Samples are aligned in order of mutation density magnitude per 1-megabase window.

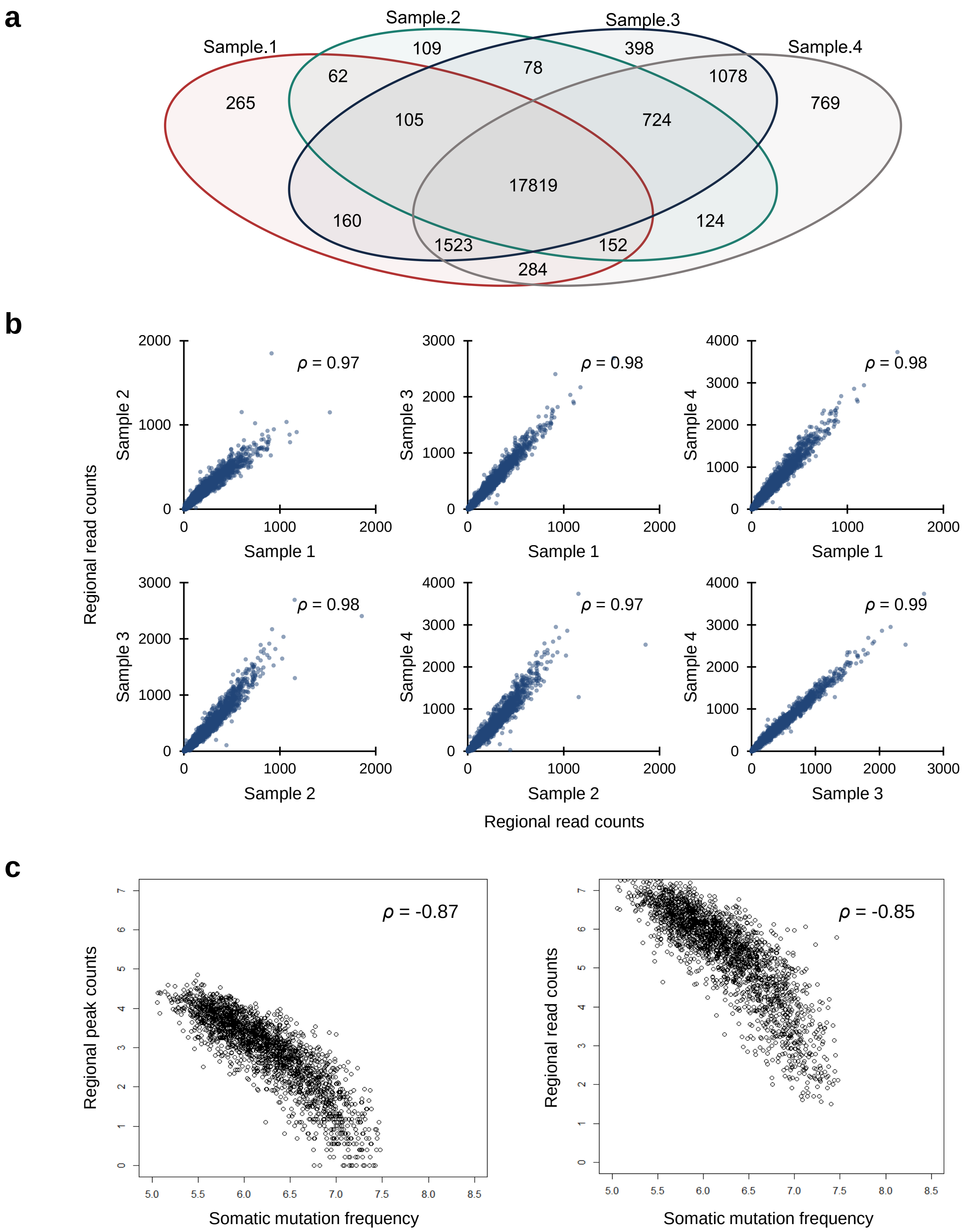


**Supplemental figure 4**. (**a**) Venn diagram of overlapping genes among individual ATAC-seq samples. (**b**) Correlation plots among the ATAC-seq samples using regional pileup counts per 1-megabase. Spearman's rank correlations ($\rho$) are shown on each plot. (**c**) Spearman correlations between the regional somatic mutation density of lung adenocarcinoma samples at the aggregate level and ATAC-seq input features derived either from the regional peak counts or pileup counts. Spearman's rank correlations ($\rho$) are shown on each plot.

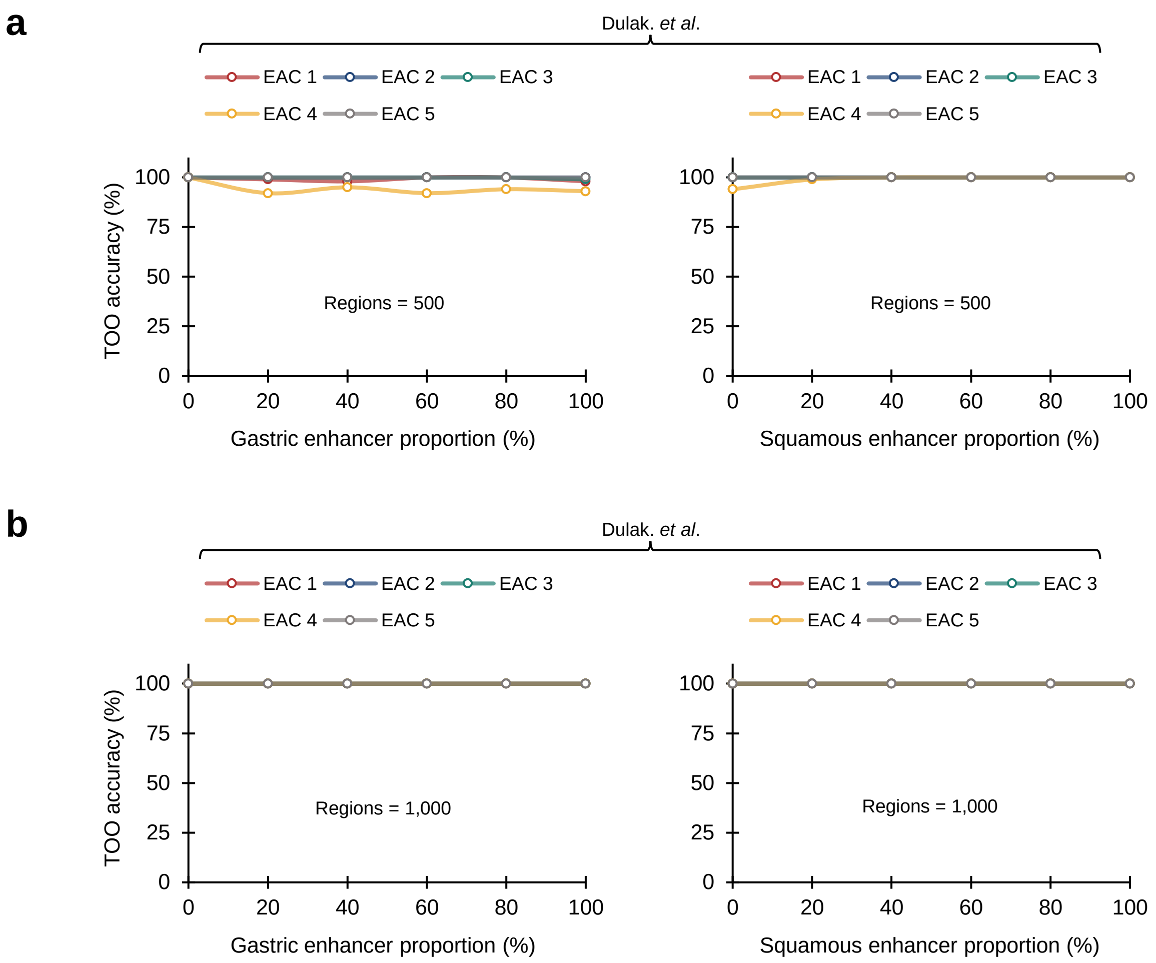


**Supplemental figure 5.** Region selection analysis with respect to the portion of gastric / squamous specific enhancer containing regions for EACs at individual sample level. Enhancer inclusion ratio was varying from 0 to 100%, making up to (**a**) 500 and (**b**) 1,000 regions.

**Supplemental table 1.**
*Speed comparison between COOBoostR and random-forest algorithm.*

| Cancer type | n | Inspection type | Average time (sec) | Min time (sec) | Max time (sec) |
|---|---|---|---|---|---|
| Colorectal cancer | 9 | Random forest 1R | 316,777 | 303,240 | 335,601 |
| | | COOBoostR 1R* | 8 | 6 | 11 |
| | | COOBoostR 100R | 761 | 577 | 1,130 |
| Esophageal adenocarcinoma Ross-Innes. *et al.* | 23 | Random forest 1R | 447,082 | 409,602 | 475,357 |
| | | COOBoostR 1R* | 16 | 10 | 26 |
| | | COOBoostR 100R | 1,588 | 1,047 | 2,615 |
| Liver cancer | 64 | Random forest 1R | 456,407 | 370,398 | 720,583 |
| | | COOBoostR 1R* | 7 | 4 | 15 |
| | | COOBoostR 100R | 707 | 383 | 1,484 |

* COOBoostR 1R values were estimated from COOBoostR 100R investigation.

**Supplemental table 2.**
*Peak and pileup statistics of ATAC-seq samples*

| Sample ID | Total peaks | Average peak length | Average pileup count |
|---|---|---|---|
| ATAC-seq sample 1 | 63,308 | 149 | 12 |
| ATAC-seq sample 2 | 61,035 | 152 | 13 |
| ATAC-seq sample 3 | 89,588 | 159 | 15 |
| ATAC-seq sample 4 | 104,916 | 159 | 17 |
